# Lack of association between Toll Like Receptor-2 & Toll Like Receptor-4 Gene Polymorphisms and Iranian Asthmatics risk or features

**DOI:** 10.1101/004382

**Authors:** Hamid Bahrami, Saeed Daneshmandi, Hasan Heidarnazhad, Ali Akbar Pourfathollah

**Affiliations:** Dept of Immunology, Tarbiat Modares University, Faculty of Medical Sciences, Tehran, Iran; TB and Lung disease Research Cancer, NRITLD, Masih Daneshvari Hospital, Shahid Beheshti University of Medical Sciences, Tehran, Iran

**Keywords:** Asthma, TLR2, TLR4, Polymorphism, RFLP-PCR

## Abstract

**Background:** Asthma as chronic inflammatory airway disease is considered to be the most common chronic disease that is involving genetic and environmental factors. Toll like receptors (TLRs) and other inflammatory mediators are important in modulation of inflammation. In this study we evaluated the role of TLR2 Arg753Gln and TLR4 Asp299Gly polymorphisms in the asthma susceptibility, progress, control levels and lung functions in Iranian subjects.

**Methods:** On 99 asthmatic patients and 120 normal subjects, TLR2 Arg753Gln and TLR4 Asp299Gly polymorphism were evaluated by PCR-RFLP method recruiting Msp1 and Nco1 restriction enzymes, respectively. IgE serum levels by ELISA technique were determined and asthma diagnosis, treatment and control levels were considered using standard schemes and criteria.

**Results:** Our results indicated that the genotype and allele frequencies of the TLR2 Arg753Gln and TLR4 Asp299Gly polymorphisms were not significantly different between control subjects and asthmatics (p > 0.05) or even in asthma features such as IgE levels, asthma history and pulmonary factors (p > 0.05).

**Conclusions:** Meanwhile some previous studies indicated TLRs and their polymorphisms role in asthma incidence and features, our data demonstrated that TLR2 Arg753Gln and TLR4 Asp299Gly gene variants were not risk factor of asthma or its features in Iranian patients. Genetic complexity, ethnicity, influence of other genes or polymorphisms may overcome these polymorphisms in our asthmatics.

## Introduction

Asthma is a complex and chronic disease in which allergen-induced inflammatory processes in the airways contribute to the development of symptoms, such as wheezing, cough, dyspnea and breathlessness [1]. Asthma is considered to be the most common chronic disease and the leading cause of hospitalization in schoolchildren [2, 3]. Control of asthma and response to medication is different in patients, and different asthmatics show various levels of asthma severity and progress which would depend on multiple factors especially genetic composition of patients. The need for management of medication and control of asthma make us improve our knowledge about pathogenesis of asthma and role of different elements that contribute to airway inflammation or influence signs and symptoms [4, 5]. Although in recent years there have been advances in pathophysiology of bronchial asthma, its cause is still unknown. Asthma is believed to be a complex disorder involving genetic and environmental factors [6]. Many allergens and environmental microorganisms have pattern molecules on their surfaces, and these molecules interact with pattern-recognition receptors, which are part of the innate immune system. Among the known pattern-recognition receptors for microbial products, toll-like receptors (TLRs) are an evolutionarily conserved group of molecules expressed in antigen-presenting cells and epithelial cells [7]. TLRs recognize microbial patterns and play a crucial role in linking innate and adaptive immunity inducing a proinflammatory immune response that may counterbalance allergic diathesis. They initiate intracellular signaling pathways, bind to downstream protein kinases, induce activation of transcription factors, and initiate transcription of inflammatory cytokines and other host response elements [8]. Among TLRs, TLR4 recognizes lipopolysaccharide, a cell wall component of gram-negative bacteria. In contrast, TLR2 recognizes a wide spectrum of PAMPs such as membrane components of Gram-positive and -negative bacterial cell wall, mycoplasma, mycobacteria, yeast and parasites [9]. A study showed the alterations in intestinal microflora balance promoted the maturity of DCs and raised the expressions of TLR2 and TLR4 on DCs in allergic mouse lung [10]. TLR2 signaling has been shown to induce Treg cell expansion accompanied by a loss of suppressive activity *in vitro* and *in vivo* [11]. A study mentioned protective effects of TLR2 polymorphisms on lung function among workers in swine operations and the possibility of its protective role in airway disease in individuals exposed to gram-positive organisms in the inhaled airborne dust [12]. In TLR4 deficient mice allergen-induced eczema were exacerbated and also Increased allergen-induced skin levels of innate (IL-1β, TNF-α, and CXCL2) and Th17 genes (IL-17A and IL-17F) were observed in TLR4-deficient mice compared with wild-type mice [13]. All of these findings support role of TLRs in asthma susceptibility or asthma features as they determine type, severity and outcomes of asthma pathogenesis. On the other hand, the amounts of TLR synthesis in the cells or their functions are determined by some polymorphisms in their genes. As a result, polymorphisms in TLRs genes might determine susceptibility or progress of disease and degree of asthma control. With regards to the proposed role TLRs in asthma, in this study we analyzed the genetic variants of TLR2 Arg753Gln, and TLR4 Asp299Gly in Iranian asthmatic patients to evaluate role of these polymorphisms in asthma susceptibility, progress, control and lung functions.

## Materials and methods

### Study populations

In this study, 99 unrelated adult patients (28 male and 71 female and mean±SD age of 44.15 ± 13.06 years), whose asthma was defined according to the criteria of the Global Initiative for Asthma (GINA) [14] were enrolled. Clinical history, physical examination, and pulmonary function test (PFT) in a standard fashion were assessed for all subjects. Asthmatics have had treatment in a standard scheme as inhaled corticosteroids and/or bronchodilator when necessary, and the asthmatic patients were subdivided into two control groups based on American Thoracic Society criteria [15] as Asthma Control Tests (ACT). Smoking history of more than 10 pack-years, presence of parasitic infection, and pregnancy or breastfeeding were exclusion criteria. A total of 120 healthy volunteers were recruited from the general population. Controls had to meet the following criteria: good health status and matched with the cases for age, gender, and area of residence, no respiratory symptoms or history of asthma and allergy. The study protocol was approved by the ethics committee at our institution, and written informed consent was obtained from all participants.

### Total serum IgE measurements

Serum was separated from 5 ml of patient and normal subjects’ blood, and total serum IgE levels were measured using the ELISA kit (Genesis Diagnostics, UK) according to the manufacturer’s instructions.

### DNA preparation

Genomic DNA was extracted from peripheral blood using a DNG^plus^ extractor WB kit (Cinagen, Iran) according to the manufacturer’s instructions.

### Determination of TLR2 Arg753Gln and TLR4 Asp299Gly gene polymorphisms

Polymerase chain reaction-restriction fragment length polymorphism (PCR-RFLP) method was used for evaluation of TLR2 Arg753Gln, and TLR4 Asp299Gly polymorphism. PCR steps were performed using a thermal cycler (Techne, Genius, UK). PCR conditions, PCR cycles and primers are summarized in Table 1 and 2. In brief, PCR materials were mixed according to table 1 and 2 and tubes passed thermal cycles as summarized in table 1. After conformation of single bands of PCR product in agarose electrophoresis, restriction enzymes were affected. In a final volume of 25 μl PCR products were digested by Msp1 and Nco1 (Fermentas thermo scientific, USA) restriction enzymes on TLR2 and TLR4 products, respectively and then digestions were monitored by agarose gel electrophoresis and ethidium bromide staining.

**Table 1:** PCR materials and cycles.

| Locus | PCR conditions |
| --- | --- |
| TLR2 | 35 cycle: 94 °C 30s, 58 °C 50s, 72 °C 40s |
|  | 50ng DNA, 200μmol dNTPs, 0.7mM MgCl<sub>2</sub> |
| TLR4 | 35 cycle: 94 °C 20s, 61 °C 50s, 72 °C 50s |
|  | 50ng DNA, 200μmol dNTPs, 2mM MgCl<sub>2</sub> |

**Table 2:**
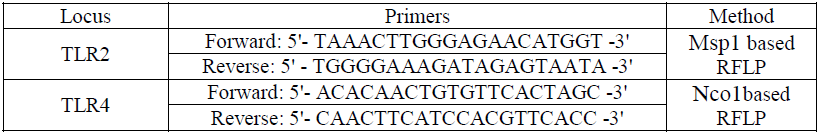
Cytokine and internal control for PCR-RFLP primers.

### Statistical analysis

Allele and genotype frequencies were calculated in patient and control subjects by direct gene counting. Statistical evaluation was carried out using the Statistical package for the Social sciences (SPSS) version 15. The statistical significance of the difference was tested by a χ^2^ analysis with one difference or by the two-tailed Fisher’s exact test when the criteria for the χ^2^ analysis were not fulfilled. The associations between genotypes and risk of asthma were estimated by computing the odds ratio (OR) and its 95% confidence interval (CI). For analysis of IgE and respiratory factors, differences in various gene variants the one way analysis variance test (ANOVA) and t-test were used. P-Values <0.05 were considered statistically significant. An exact test was used to evaluate deviations from expected Hardy-Weinberg genotypic proportions.

## Results

The results of TLR2 Arg753Gln, and TLR4 Asp299Gly gene polymorphisms in asthmatic and normal subjects (Table 3), two levels of asthma controls (Table 4), sex, allergy history and familial history of asthma (Table 5) and serum IgE and respiratory factors in asthma patients (Table 6) are shown. Our results analysis didn’t show any statistically significant difference between TLR2 Arg753Gln and TLR4 Asp299Gly genotypes and alleles of asthmatics and normal subjects (P>0.05). Other features of asthma as asthma control levels, serum IgE, history and pulmonary factors also were not different in TLR2 Arg753Gln and TLR4 Asp299Gly different variants (P>0.05).

**Table 3:** Results of TLR2 Arg753Gln and TLR4 Asp299Gly SNPs determined in asthmatics and normal subjects.

| Gene | Genotype | Asthma<br>%(N) | Normal<br>%(N) | P value | OR | %95CI |
| --- | --- | --- | --- | --- | --- | --- |
| TLR2 | GG | %92.9(92) | %94.2(113) | 0.709 | 0.814 | 0.276-2.405 |
|  | GA+AA | %7.1(7) | %5.8(7) |  |  |  |
|  | G allele | %96.5(191) | %97.1(233) | 0.714 | 0.820 | 0.283-2.378 |
|  | A allele | %3.5(7) | %2.9(7) |  |  |  |
| TLR4 | AA | %85.9(85) | %86.7(104) | 0.863 | 0.934 | 0.431-2.022 |
|  | AG+GG | %14.1(14) | %13.3(16) |  |  |  |
|  | A allele | %92.9(184) | %93.3(224) | 0.868 | 0.939 | 0.446-1.974 |
|  | G allele | %7.1(14) | %6.7(16) |  |  |  |

N, absolute number; CI, Confidence Interval; OR, Odds ratio.

**Table 4:**
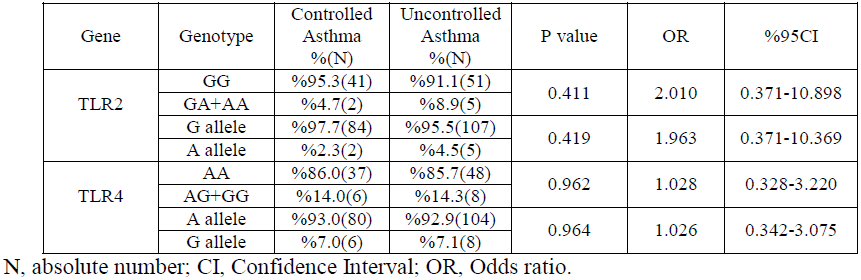
Results of TLR2 Arg753Gln and TLR4 Asp299Gly SNPs distribution in asthma control levels.

**Table 5:** Distribution and P values for TLR2 Arg753Gln and TLR4 Asp299Gly variants in sex, allergy history and familial history of asthma groups.

| Gene | Genotype | Distribution (N) |  |  | P value |  |  |
| --- | --- | --- | --- | --- | --- | --- | --- |
|  |  | Sex M/F | Allergy P/N | History P/N | Sex | Allergy | History |
| TLR2 | GG | 26/66 | 63/29 | 44/48 | 0.986 | 0.077 | 0.800 |
|  | GA+AA | 2/5 | 7/0 | 3/4 |  |  |  |
|  | G allele | 54/137 | 133/58 | 91/100 | 0.986 | 0.077 | 0.803 |
|  | A allele | 2/5 | 7/0 | 3/4 |  |  |  |
| TLR4 | AA | 25/60 | 59/26 | 39/46 | 0.539 | 0.485 | 0.434 |
|  | AG+GG | 3/1 | 11/3 | 8/6 |  |  |  |
|  | A allele | 53/131 | 129/55 | 86/95 | 0.555 | 0.502 | 0.452 |
|  | G allele | 3/11 | 11/3 | 8/6 |  |  |  |

M: Male; F: Female; P: Positive; N: Negative.

**Table 6:** IgE concentrations and P values for association between TLR2 Arg753Gln and TLR4 Asp299Gly polymorphisms and serum IgE and respiratory factors of asthma patients.

| Gene | Genotype | IgE Cons.<br>(IU/ml) | P value |  |  |  |  |  |  |
| --- | --- | --- | --- | --- | --- | --- | --- | --- | --- |
|  |  |  | IgE | Age | FEV1<br>(%P) | FVC<br>(%P) | FEV1/FVC<br>(%P) | FEF<br>25-75%<br>(%P) | PEF<br>(%P) |
| TLR-2 | GG | 113.84±251.38 | 0.258 | 0.658 | 0.928 | 0.731 | 0.663 | 0.282 | 0.436 |
|  | GA+AA | 241.61±564.32 |  |  |  |  |  |  |  |
|  | G allele | 119.16±268.71 | 0.265 | 0.662 | 0.929 | 0.733 | 0.256 | 0.665 | 0.440 |
|  | A allele | 241.61±564.32 |  |  |  |  |  |  |  |
| TLR-4 | AA | 117.37±257.65 | 0.584 | 0.371 | 0.158 | 0.295 | 0.948 | 0.427 | 0.238 |
|  | AG+GG | 116.20±433.55 |  |  |  |  |  |  |  |
|  | A allele | 120.94±271.91 | 0.596 | 0.388 | 0.174 | 0.315 | 0.949 | 0.440 | 0.257 |
|  | G allele | 166.20±433.55 |  |  |  |  |  |  |  |

Forced Expiratory Volume in 1 Second, FVC: Forced Vital Capacity, PEF: Peak Expiratory Flow, FEF25-75: Forced Expiratory Flow 25-75%.

## Discussion

Asthma is a multifactor chronic inflammatory disorder of the airways and a variety of genetic and environmental factors contribute to its pathogenesis. Immune and inflammatory elements are important factors in induction, progress and clinical outcomes of asthma [6]. Molecules and elements that are linker of environmental factors effects in these inflammatory responses and then their genetic background will be critical factors in asthma etiology or features. TLRs are pattern recognition receptors, are highly polymorphic, and play an important role in both innate and adaptive immunity [9, 16]. TLR2 mainly responds to cell wall structure components from gram-positive bacteria, such as peptidoglycan, and TLR4 mainly recognizes microbial membrane components from gram-negative bacteria, such as lipopolysaccharide (LPS) orendotoxin [9, 17]. TLRs polymorphisms may play role in the balance between Th1 and Th2 responses, and increased susceptibility to allergic airway diseases. In a study to quantify messenger RNA (mRNA) and protein expression of TLR2, TLR3, and TLR4 in the nasal mucosa of patients with seasonal allergic rhinitis before and after challenge with relevant pollens identified, protein expression for all three TLRs was demonstrated to increase in patients with rhinitis after challenge. This study raises the possibility that TLRs polymorphisms may have a role in the development of allergic airway inflammation [18]. TLR2 is encoded at 4q32 and TLR4 located at 9q32-33. TLR4 polymorphisms have been associated with reduced risk of allergic rhinitis, atopy, and airway responsiveness in several studies [19, 20] The reduced immune responses of TLR2 polymorphisms have also been investigated in experimental studies using human cells or animal models [21, 22]. All of these data support TLRs molecules and also their polymorphisms roles in asthma features. With regard to these findings we evaluated two main polymorphisms as TLR2 Arg753Gln, and TLR4 Asp299Gly in Iranian asthmatic patients and then analyzed correlation of these variants with serum IgE levels, respiratory factors, patient’s allergy and familial history and also standard control levels of asthma in these patients. Analysis of our results didn’t show any positive correlation of TLR2 Arg753Gln, and TLR4 Asp299Gly polymorphisms with asthma risk, IgE levels, lung functions, control level or even familial allergy/asthma history. For explanation of these results we looked up at some previous studies.

A study on Chinese asthmatics analyzed four SNP in TLR4 gene resulted polymorphisms in TLR4 gene are associated with asthma severity but not susceptibility [23]. These researchers in another paper on the same patients reported positive correlation TLR2/rs7656411 TT variant and risk of asthma [24]. In a study single nucleotide polymorphisms (SNPs) in the TLR2 (4 SNPs) and TLR4 genes (9 SNPs) were genotyped in asthmatic children. In these subjects Two TLR2 SNPs and four TLR4 SNPs significantly modified the effect of air pollution on the prevalence of doctor-diagnosed asthma [25]. A study reported protective effects of TLR2-16933T/A polymorphisms on lung function among workers in swine operations while in these subjects there were no significant differences between Asp299Gly and Thr399Ile polymorphisms in the TLR4 gene and lung function values [26]. In another study on Danish asthmatic farmers three CD14 SNPs, three TLR2 SNPs (-16934 A/T, Pro631His C/A and Arg753Gln C/T), and two TLR4 SNPs (Asp299Gly A/G and Thr399Ile C/T) were evaluated and their results indicated no associations between CD14, TLR2, or TLR4 genotypes and new-onset asthma [27]. In another study on South Korean allergic rhinitis patients there were not a significant correlation between TLR2 Arg753Gln, and TLR4 Asp299Gly polymorphisms and allergic rhinitis risk and also serum IgE levels [28]. Eder et al found that children with a Tallele of TLR2-16934 (rs4696480) growing up on a farm were less likely to have a diagnosis of asthma [29] Werner et al showed that two non-synonymous SNPs of the TLR4 gene (rs4986790 and rs4986791) modified the effect of endotoxin exposure on asthma in adults [30]. A study on Egyptian asthmatics demonstrated a lack of association of TLR2 Arg753Gln and TLR4 Asp299Gly polymorphisms with asthma and allergic rhinitis but suggested significant association between these genetic variants and the disease severity. Another study were found that a polymorphism as TLR2/–1693 is strongly associated with the frequency of asthma and allergies in children of European farmers. In these subjects two other TLR2 SNPs and five polymorphism of TLR4 didn’t have correlation with asthma [29]. In a quick look to the previous studies and with attention to probable role of TLRs in asthma we see several studies that have mentioned polymorphisms or other SNPs in TLR2 and TLR4 genes are involved in asthma and meanwhile there are also several studies that have showed no such association. These controversies may be due to several explanation such as: genetic complexity and/or nature of asthma pathogenesis and pathophysiology with involvement of various factors including genetic and environmental factors, ethnicity and different genetic background of studied subjects, overcame of other molecules and their genetic variations on TLRs functions and even effectiveness of other polymorphisms in TLR2 and TLR4 genes more than two TLR2 Arg753Gln and TLR4 Asp299Gly studied polymorphisms in Iranian asthmatic patients. We would mention that in previous study in almost the same asthmatics, we showed effectiveness of cytokine polymorphism in asthma susceptibility and its features in Iranian asthmatic patients [32]. our results in this study denyed to indicate TLR2 Arg753Gln and TLR4 Asp299Gly polymorphisms as a risk factor for asthma in Iranian patients. These studies SNPs also didn’t affect asthma features as IgE levels, lung functions, control level or even familial allergy/asthma history. With regard to positive correlations in some of other studies and supposed roles of TLR2 and TLR4 in modification of immune and inflammatory responses further studies on other SNPs or other involved may be beneficial for clarification of controversies in multifactor asthma disease.

## Acknowledgement

The authors are grateful to the Department of Immunology of Tarbiat Modares University for financial support.

## Declaration of interest

The authors report no conflicts of interest. The authors alone are responsible for the content and writing of article.

